# Strongly asymmetric hybridization barriers shape the origin of a new polyploid species and its hybrid ancestor

**DOI:** 10.1101/030932

**Authors:** Marío Vallejo-Marin, Arielle M. Cooley, Michelle Lee Yue Qi, Madison Folmer, Michael R. McKain, Joshua R. Puzey

## Abstract

- *Premise of the study:* Hybridization between diploids and tetraploids can lead to new allopolyploid species, often via a triploid intermediate. Viable triploids are often produced asymmetrically, with greater success observed for “maternal-excess” crosses where the mother has a higher ploidy than the father. Here we investigate the evolutionary origins of *Mimulus peregrinus*, an allohexaploid recently derived from the triploid *M. x robertsii*, to determine whether reproductive asymmetry has shaped the formation of this new species.
- *Methods:* We used reciprocal crosses between the diploid (*M. guttatus*) and tetraploid (*M. luteus*) progenitors to determine the viability of triploid *M. x robertsii* hybrids resulting from paternal-versus maternal-excess crosses. To investigate whether experimental results predict patterns seen in the field, we performed parentage analyses comparing natural populations of *M. peregrinus* to its diploid, tetraploid, and triploid progenitors. Organellar sequences obtained from pre-existing genomic data, supplemented with additional genotyping was used to establish the maternal ancestry of multiple *M. peregrinus* and *M. x robertsii* populations.
- *Key results:* We find strong evidence for asymmetric origins of *M. peregrinus*, but opposite to the common pattern, with paternal-excess crosses significantly more successful than maternal-excess crosses. These results successfully predicted hybrid formation in nature: 111 of 114 *M. x robertsii* individuals, and 27 of 27 *M. peregrinus*, had an *M. guttatus* maternal haplotype.
- *Conclusion:* This study, which includes the first *Mimulus* chloroplast genome assembly, demonstrates the utility of parentage analysis through genome skimming. We highlight the benefits of complementing genomic analyses with experimental approaches to understand asymmetry in allopolyploid speciation.

## INTRODUCTION

The extraordinary abundance of polyploid lineages among flowering plants (Otto and Whitton, 2000; Soltis et al., 2014; Barker et al., 2015) has led to great interest in understanding the processes of polyploid formation and establishment in the wild. Polyploids (individuals carrying more than two sets of chromosomes) can originate through a variety of pathways. These include the somatic doubling of chromosomes in meristematic tissue during mitosis and, perhaps more commonly, unreduced gametes (Ramsey and Schemske, 1998; Mallet, 2007). When the formation of polyploids involves fusion of gametes with different ploidy levels (e.g., reduced and unreduced gametes, or between a diploid and a tetraploid), an individual with an unbalanced ploidy level (e.g., a triploid) will be formed (Levin, 2002). In most cases, these odd-ploidy individuals are a transient phase in polyploid formation, leading to the eventual production of higher ploidy, stable lineages such as tetraploids and hexaploids (Levin, 2002).

Odd-ploidy levels represent a significant challenge to overcome in the formation of polyploids, as inter-ploidy crosses often display strong post-zygotic reproductive barriers, severely reducing hybrid viability and fertility (Ramsey and Schemske, 1998; Levin, 2002; Comai, 2005; Köhler, Scheid, and Erilova, 2010). The term “triploid block” has been coined to describe the regular observation that inter-ploidy crosses, specifically between diploids and tetraploids, often result in early hybrid inviability and sterility (Köhler, Scheid, and Erilova, 2010). However, evidence from both natural and experimental systems has shown that interploidy crosses can sometimes produce odd-ploidy offspring that are viable (Burton and Husband, 2000; Husband, 2004; Stace, Preston, and Pearman, 2015).

The ability to overcome the triploid block can be asymmetric, with offspring viability dependent on whether the maternal or paternal parent had the higher ploidy level (Köhler, Scheid, and Erilova, 2010). Asymmetric hybridization has considerable potential for influencing population genetic processes. It generates a predictable bias in cytoplasmic inheritance, and in some cases can promote unidirectional introgression (Field et al., 2011). Repeated backcrossing between an asymmetrically-produced hybrid and its paternal pollen source may lead to “cytoplasmic capture,” in which the mitochondria and chloroplasts of the maternal progenitor are combined with the nuclear genome of the paternal progenitor (Petit et al., 2004). Asymmetric introgression can have conservation implications, for example through genetic swamping or competitive displacement of the rarer progenitor species (Rhymer and Simberloff, 1996).

Asymmetric hybridization can be explained by a variety of mechanisms, including cyto-nuclear incompatibilities (Tiffin, Olson, and Moyle, 2001), or imbalances in either the ploidy ratio of endosperm:embryo (Watkins, 1932; Burton and Husband, 2000) or between maternal:paternal genomes (Haig and Westoby, 1989; Köhler, Scheid, and Erilova, 2010). Barriers to inter-ploidy mating within and between species, including triploid block, have been studied using controlled crosses in a variety of natural and artificially produced diploids and polyploids (Ramsey and Schemske, 1998; Husband, 2004; Köhler, Scheid, and Erilova, 2010; Scott, Tratt, and Bolbol, 2013). Earlier surveys of interploidy crossing data determined that maternal-excess crosses— those in which the maternal parent has a higher ploidy level than the paternal one—are consistently more successful than paternal-excess crosses at producing viable triploid seeds (Stebbins, 1957; Ramsey and Schemske, 1998). However, a number of counter-examples, in which paternal-excess crosses are more successful, have been recently discovered (Table 1; Appendix S1).

**Table 1.**
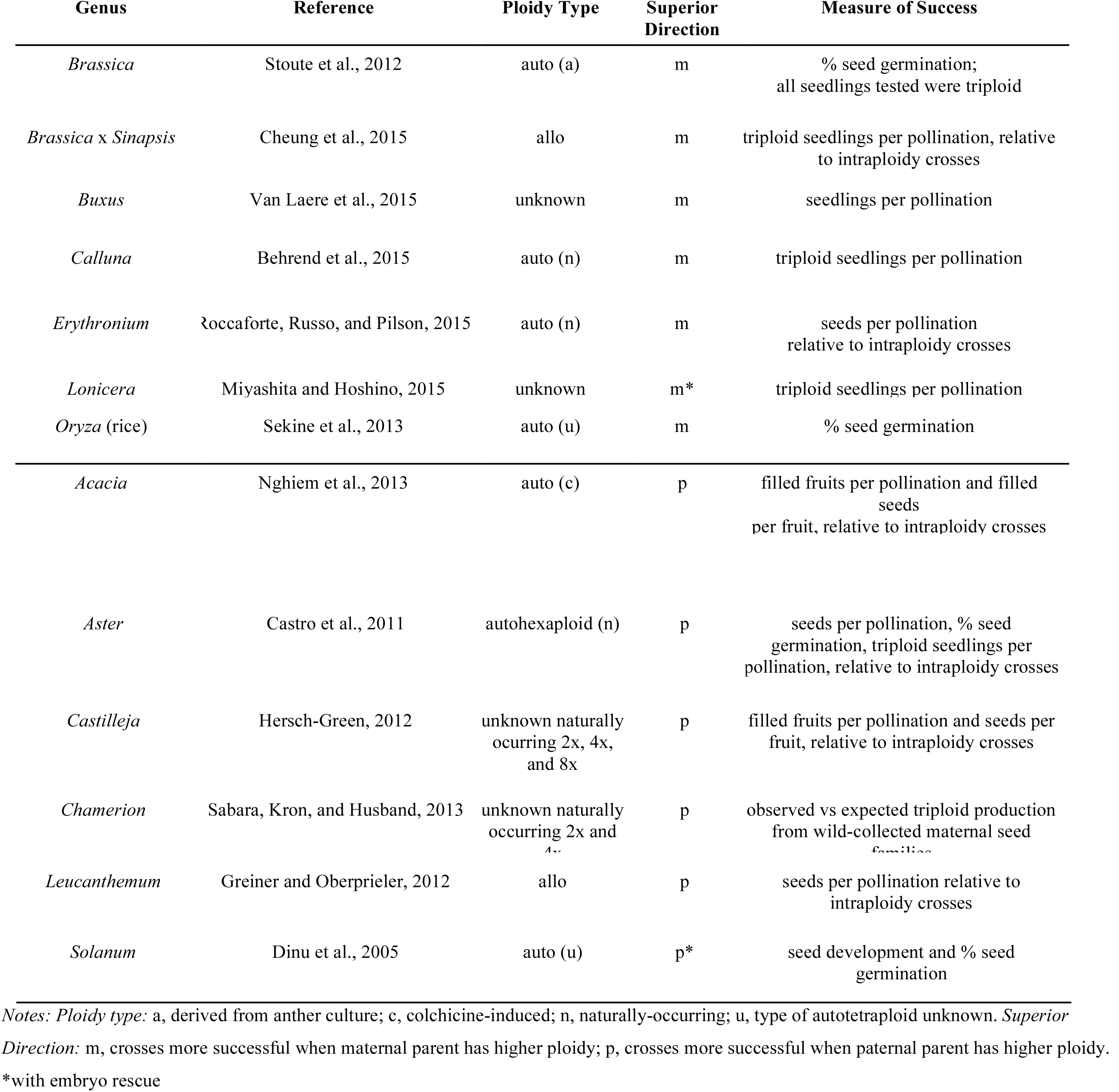
Recent (1998-2015) reports of asymmetric crossing success following interploidy crosses. Some of the studies listed below verified that the seeds or seedlings produced were triploid; others did not and could potentially include diploid, tetraploid, or aneuploid offspring in their measure of success.

Asymmetric reproductive barriers during interploidy hybridization can be identified experimentally by performing reciprocal crosses (e.g. Husband, 2004), but discovering whether asymmetries observed in the lab translate to patterns in nature, requires tracing the maternal *versus* paternal ancestry of wild populations. Ancestry analysis of this kind requires uniparentally inherited genetic markers, such as mitochondria or chloroplast that contain enough genetic differences between parental taxa. The rapid growth of next generation sequencing projects offers an extraordinary opportunity to generate (sometimes inadvertently) exactly the type of data necessary to elucidate the maternal ancestry of wild populations at a genomic scale. Because cytoplasmic genomes (mitochondria and chloroplast) are present at a much higher number per cell than the nuclear genome, targeted and non-targeted sequencing often results in incidental high coverage of cytoplasmic genomes, as well as of repetitive elements in the nuclear genome (Dodsworth, 2015). Thus mitochondrial and chloroplast information can be “skimmed” from the larger collection of sequences obtained in whole-genome sequencing projects (Straub et al., 2012), or from targeted sequencing approaches (Weitemier et al., 2014), and used for parentage analyses. These resource-efficient approaches not only rescue information that would otherwise be discarded, but also provide effective means for addressing longstanding questions about the ancestries of naturally-occurring polyploid lineages.

A new system for investigating allopolyploid speciation, using the complementary approaches of experimental manipulation and next-generation sequencing, is the complex of naturalized *Mimulus* L. (Phrymaceae) species in Great Britain and Ireland. Hybridization between introduced populations of the North American diploid *M. guttatus* (2n = 2x = 28) and the South American tetraploid *M. luteus* (2n = 4x = 60-62) have given rise to *M. x robertsii*, a triploid (2n = 3x = 44-46), sexually sterile, but widely naturalized taxon distributed across the British Isles (Roberts, 1964; Parker, 1975; Silverside, 1990; Vallejo-Marín et al., 2015). By means of rapid vegetative propagation, *M. x. robertsii* has become common and is now found in about 40% of British *Mimulus* populations (Vallejo-Marín and Lye, 2013). The reproductive asymmetry typical of interploidy matings has been reported for *M. guttatus* x *M. luteus* hybrids, but in the less common direction: triploids have been successfully produced only by paternal-excess crosses, with tetraploid *M. luteus* as the pollen donor (Roberts, 1964; Parker, 1975). These early studies suggest that *M. x robertsii* is likely to have arisen in the same way, from a *M. guttatus* mother and *M. luteus* father. *Mimulus x robertsii* has given rise at least twice to a sexually fertile allohexaploid species, *M. peregrinus* Vallejo-Marín (2*n* = 6*x* = 92), through genome doubling (Vallejo-Marín, 2012; Vallejo-Marín et al., 2015). This speciation event is quite recent, as both progenitors were introduced to the United Kingdom (UK) in the last 200 years, and the earliest records of hybrid populations date back approximately 140 years (Preston, Pearman, and Dines, 2002; Vallejo-Marín et al., 2015). Given that *M. peregrinus* evolved from *M. x robertsii*, one would expect that the maternal ancestor of *M. peregrinus* is also *M. guttatus*.

Here we investigate the occurrence and consequences of inter-ploidy reproductive barriers in polyploid *Mimulus* (*M. x robertsii* and *M. peregrinus*) produced by hybridization between diploid (*M. guttatus*) and tetraploid taxa (*M. luteus*). We complement classic crossing experiments with analyses of the parentage of natural hybrid populations using genome-skimming of chloroplast and mitochondrial genomes and single-marker genotyping. Inter-ploidy hybrids in *Mimulus* are ideally suited for this purpose due to their experimental tractability (Wu et al., 2008) as well as the growing genomic resources available for this genus, including whole genome sequences for both nuclear and mitochondrial genomes for *M. guttatus* (Mower et al., 2012; Hellsten et al., 2013), ongoing work on the *M. luteus* genome (Edger et al., in prep), and recent whole-genome and targeted sequence data for native and introduced *Mimulus* populations (Puzey and Vallejo-Marín, 2014; Vallejo-Marín et al., 2015). We experimentally evaluate the success of reciprocal crosses between *M. guttatus* and *M. luteus* to confirm previous observations of triploid block asymmetry, and compare the phenotype of hybrids produced in reciprocal directions. We use a combination of genome skimming and genotyping of natural populations to evaluate the hypothesis that reproductive asymmetry observed between *M. guttatus* and *M. luteus* violates the trend (Ramsey and Schemske, 1998) of triploid formation via maternal-excess crosses. Our study addresses the following specific questions: (1) What is the effect of cross direction on the phenotype and viability of hybrids? (2) Are naturalized triploid hybrids between *M. guttatus* and *M. luteus* (i.e., *M. x robertsii*) asymmetrically produced, and does the asymmetry match the pattern observed in experimental hybridizations? (3) Who is the maternal parent of hexaploid *M. peregrinus*, and does this fit the pattern seen for experimental crosses and naturalized *M. x robertsii*?

## MATERIAL AND METHODS

***Study system***—The introduction of *Mimulus* into Europe in the 1800’s resulted in hybridization between closely related, but previously geographically isolated taxa. Hybridization between taxa in the North American *M. guttatus* DC. and South American *M. luteus* L. species aggregates has yielded a taxonomically complex array of hybrids that establish naturalized populations in the British Isles (Roberts, 1964; Silverside, 1998; Stace, Preston, and Pearman, 2015). The *M. luteus* group includes a number of interfertile taxa: *M. luteus* var. *luteus, M. luteus* var. *variegatus, M. naiandinus, M. cupreus*, and *M. depressus* (Grant, 1924; Watson, 1989; von Bohlen, 1995; Cooley and Willis, 2009), some of which have become naturalized in the British Isles (Stace, 2010). Extant populations of *M. luteus* in the British Isles include highly polymorphic individuals, which may have resulted from hybridization events between different naturalized varieties. Here we focus on interspecific hybrids between *M. guttatus* and different varieties of *M. luteus s.l*., excluding hybrids with another closely related South American taxon, *M. cupreus*, which are rarer and easily distinguished (Stace, 2010). Hybrids between *M. guttatus* and *M. luteus s.l*. receive different taxonomic names, mostly based on color patterns of the corolla (Stace, 2010). For simplicity, hereafter we refer to these hybrids as *M. x robertsii*, and the interested reader is referred to Silverside (1998) for a more detailed morphological classification of hybrid types.

Hybridization between diploid *M. guttatus* (2n= 2x = 28) and tetraploid *M. luteus* (2n = 4x = 60-62) yields highly sterile, triploid individuals (2n = 3x = 44 - 46, *M. x robertsii*). Despite their sterility (Roberts, 1964; Parker, 1975), these triploid hybrids can be vegetatively vigorous and display a strong capacity to reproduce by clonal propagation from plant fragments that root at the nodes. *M. x robertsii* has been very successful at establishing naturalized populations throughout the British Isles (Stace, Preston, and Pearman, 2015), and it can become locally abundant, forming populations of thousands of individuals. A recent genetic survey of *M. x robertsii* showed that populations of this sterile hybrid are genetically variable and composed of multiple highly heterozygous genotypes (Vallejo-Marín and Lye, 2013). Approximately 40-50% of extant *Mimulus* populations in the United Kingdom contain *M. x robertsii* (Vallejo-Marín and Lye, 2013).

***Crossing design***—To generate the plants used in this study, seeds were collected from natural populations of *M. guttatus* from Dunblane, Perthshire, Scotland (56.19°N, 3.98°W), and *M. luteus s.l*. from Coldstream, Scottish Borders, Scotland (55.65°N, 2.24°W) on 8 Aug 2009 and 10 July 2010, respectively. Individuals of *M. luteus s.l*. from Coldstream are polymorphic for the number of purple blotches present in the corolla lobes. Phenotypically, these *M. luteus s.l*. plants resemble intraspecific hybrids between *M. luteus* var. *luteus* and *M. luteus* var. *variegatus* (sometimes called *M. smithii*). From each population, we randomly chose six individuals collected as seeds from separate maternal plants and crossed them to generate 12 outbred lines for each species using an equal contribution design in which each individual contributed to the genome of four lines, two as the paternal parent and two as the maternal parent.

To generate interspecific hybrids, each of the 12 *M. guttatus* outbred lines were randomly paired with one of 12 *M. luteus s.l*. outbred lines. Two of these interspecific pairs were then assigned to each of six mating groups, and each of these groups was subject to a diallel cross, except selfs. To avoid half-sib matings, conspecifics in the same mating group were chosen to be unrelated. Therefore, all matings within each group are outbred and have the same inbreeding coefficient relative to the parental generation. Flowers were emasculated before conducting hand pollinations. Only 33 out of 72 possible hybrid crosses were performed due to flower availability.

The crossing design yielded individuals of four different types: (**a**) *M. guttatus* x *M. guttatus* (MG x MG); (**b**) *M. luteus s.l*. x *M. luteus s.l*. (ML x ML); and reciprocal F1 hybrid crosses of (**c**) *M. guttatus* x *M. luteus s.l*. (MG x ML), and (**d**) *M. luteus s.l*. x *M. guttatus* (ML x MG). Hereafter, crosses are represented as maternal x paternal parent. In total we obtained seeds of 33 lines of the following crossing types: MG x MG (4 lines), ML x ML (10 lines), MG x ML (10 lines), and ML x MG (9 lines). For all subsequent experiments, we randomly chose four lines of each crossing type (16 lines total; Appendix S2; see Supplemental Data with the online version of this article).

***Seed production***—To determine the number of seeds produced per fruit, between 14 to 20 days after hand pollination, 9 to 11 randomly selected fruits of each parental and F1 cross types were collected into glassine paper bags. The fruits were dissected in a stereo microscope (MZ6, Leica Microsystems, Milton Keynes, UK) at 0.63x – 4.0x magnification to remove the fruit casing and count all seeds.

***Germination rate***—To determine seed viability and germination, for the F1 crosses, 100 randomly selected seeds from each line were surface-sown in 3.5”, 370 mL, round pots (9F; Desch PlantPak, Netherlands) using a low nutrient compost (Modular Seed Growing Medium, William-Sinclair, Lincoln, UK). The pots were arranged in plastic trays in standing water (approximately 3-4cm water depth). Trays were placed in controlled environment chambers (Microclima; Snijders Scientific B.V, Tilberg, Netherlands) at the University of Stirling, and maintained at cycles of 16 light-hours (24°C) and 8 dark-hours (16°C) at 70% constant humidity. Seedling emergence was scored daily by counting the cumulative number of individuals in which the cotyledons could be observed.

***Floral and vegetative phenotyping***—To compare the phenotypes of parental and hybrid lines, a second set of seeds from the same lines were planted and germinated as described above. Individual seedlings of each line were transplanted 10 days after germination into 3.5” pots with low nutrient compost and arranged in flooded, plastic trays. Trays were placed at the University of Stirling glasshouses under natural light conditions, with 16 hours of supplemental light provided by compact-fluorescent lamps. Heating was only provided if the temperature inside the glasshouse dropped below 15°C during the day and 10°C at night. Plants were sprayed once to control for an aphid infestation, with 1.5L of diluted Provado Ultimate Bug Killer (20 mL/L) (Bayer Garden, Cambridge, UK). This experiment was carried out between 3 July and 2 October 2012. The date and plant height at the time when the first flower opened was recorded, as well as the following measurements from the first two flowers of each plant, which were taken using digital calipers: (a) corolla width (maximum distance between lateral petal lobes in frontal view); (b) corolla height (maximum distance between upper and lower petal lobes in frontal view); (c) corolla throat (size of the opening of the corolla tube taken near the center of the flower); (d) calyx length (measured at the upper sepal); (e) tube length (measured in side view); (f) pedicel length; (g) bract width (width of the subtending leaf at the node of the flower being measured); and (h) bract length (including petiole). Nectar production was measured on a subset of flowers using a 50µL calibrated glass pipette (Drummond Scientific Company, Broomall, PA, USA). The fertility of reciprocal crosses was not quantified here, but previous work (Roberts 1964), and our preliminary observations suggest that triploid hybrids are sterile regardless of crossing direction.

***Confirming the success of artificial crosses***—To further check that the individuals produced through artificial crosses were indeed derived from interspecific hybridization, we genotyped all plants used in the morphological analyses. Leaf tissue samples were collected in August-October 2012 and stored in silica gel. DNA was extracted using a modification of the CTAB protocol described in Doyle and Doyle (1990), and quantified with Nanodrop 2000 (Thermo Scientific, Wilmington, DE, USA). We genotyped putative hybrids and parents at six microsatellite loci (ATT240, AAT217, ATT267, AAT230, AAT225, AAT300; (Kelly and Willis, 1998) following the protocol described in Vallejo-Marín and Lye (2013). We used STRand 2.4.59 (Toonen and Hughes, 2001) to analyze fluorescence profiles. Based on the genotyping results, four ML x MG individuals were identified as self-pollinations of the ML maternal plant based on lack of expected heterozygosity, and excluded from subsequent analyses. While we do not have cytological data for individuals included in this experiment, we have confirmed the triploid nature of other putative triploid individuals in the same interspecific crosses using genome size estimates of nuclei stained with propidium iodide, and analyzed in a *Guava easyCyte 5* (Merck Millipore, Watford, U.K.) flow cytometer (Vallejo-Marín, unpublished).

***Data analysis***—The number of seeds produced by the four cross types (MG x MG, MG x ML, ML x MG, and ML x ML) was statistically compared with a general linear model (with normal distribution) with cross type as explanatory variable using the *glm* package in *R* ver. 3.1.2 (R Core Team, 2015). Pairwise posthoc comparisons among cross types was done using Tukey tests on the fitted model using *multcomp* (Hothorn, Bretz, and Westfall, 2008). Germination curves for each cross type were calculated as the mean of the four studied lines. The proportion of plants that flowered was analyzed using a binomial distribution (logit link) in *glm*, and posthoc pairwise comparisons among cross types done with Tukey tests in *multcomp*. Nectar content and plant height were also analyzed using *glm* with cross type as an explanatory variable, and Tukey tests performed in *multcomp*. Floral and leaf traits are presented as mean and standard error for illustration only, and no further general linear model or pairwise posthoc comparisons for cross types were done on these traits. A PCA of the 10 floral and vegetative traits shown in Table 4 was conducted using the function *princomp* on the correlation matrix.

***Chloroplast assembly***—Despite the availability of both nuclear (Hellsten et al., 2013) and mitochondrial (Mower et al., 2012) genomes for *M. guttatus*, no chloroplast genome for any *Mimulus* species had been previously published. Thus we sought to generate an assembly of the entire chloroplast genome using newly generated whole-genome sequences of an advanced inbred line of *M. luteus var. luteus* (MLll/MLl2, Table 2). MLll/MLl2 are the same inbred line but are listed separately in Table 2 as they were genotyped via different methods (WGS and sequence capture). Whole genome shotgun reads were trimmed using Trimmomatic v.0.32 (Bolger, Lohse, and Usadel, 2014) with the SLIDINGWINDOW:10:20 and MINLEN:40 parameters. Trimmed reads were mapped against the *Salvia miltiorrhiza* chloroplast genome (NC_020431.1; Qian et al., 2013) using bowtie2 v.2.1.0 (Langmead and Salzberg, 2012) with the default “very-sensitive-local parameter” set. Mapped reads were assembled using SPAdes v.3.1.0 (Bankevich et al., 2012) with k-mer sizes of 55 and 87 under the “only-assembler” option. The resulting contigs were assembled with the full trimmed data set using afin (bitbucket.org/benine/afin) with the following parameters: a stop extension value of 0.1, an initial trim of 100 base pairs to contigs, a maximum extension of 100 bp per loop, and 50 search loops. afin spans contigs by trimming contig ends, identifying matching reads to the contig ends, and extending the contigs iteratively while attempting to fuse contigs at each iteration. Ultimately, afin was able to assemble the plastome from ˜10 contigs to one. The assembled plastome was put into Sequencher 5.0.1 (Genecodes), and the boundaries of the quadripartite regions (large single copy-LSC, inverted repeat B-IRB, small single copy-SSC, and inverted repeat A-IRA) were identified. The final plastome is presented in the LSC-IRB-SSC-IRA format. The final assembly of the plastome was verified through a coverage analysis. Jellyfish v.2.1.3 (Marçais and Kingsford, 2011) was used to estimate 25-mer abundance from the cleaned reads. These abundances were used to map a 25 base pair sliding window of coverage across the assembled plastome. The coverage was found to be uniform and equal for the single copy regions (LSC=373X; SSC=328X) with the inverted repeat having ˜2x the coverage of the single copy regions (IR=756X), as expected. DOGMA (Wyman, Jansen, and Boore, 2004) was used to annotate the *Mimulus luteus* plastome. A graph of the annotated plastome structure was created using Circos v.0.66 (Krzywinski et al., 2009), as implemented in Verdant (verdant.iplantcollaborative.org).

**Table 2.**
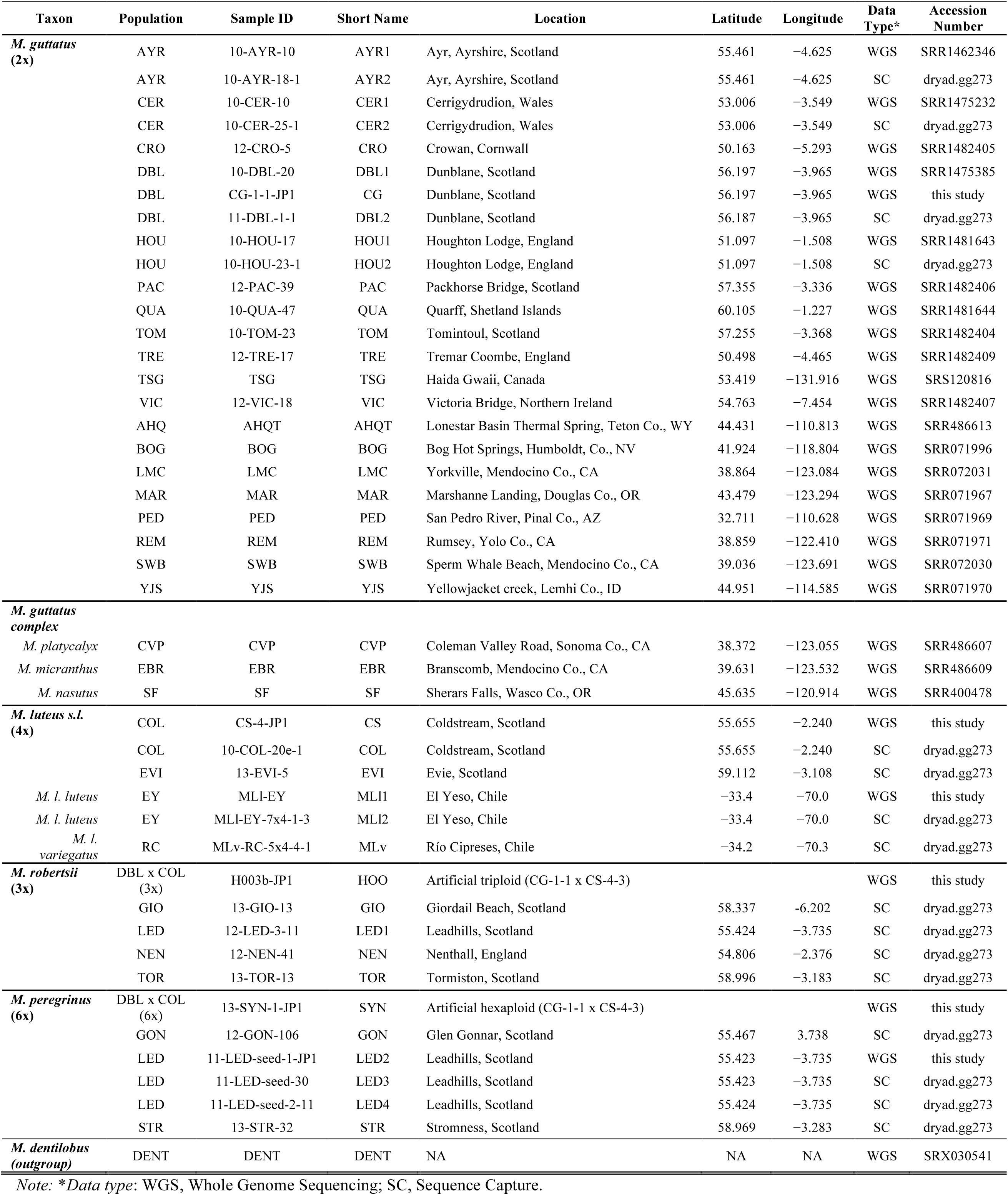
Sample names and sampling locations for *Mimulus* accessions used in this study. Data type: WGS = whole-genome sequencing; SC = sequence capture.

***Samples for genomic parentage analysis***—In order to determine the relationships between the cytoplasmic genomes of *M. guttatus* and *M. luteus*, we analyzed previously published and newly generated sequencing data of both parental taxa (Table 2). We included 24 individuals of *M. guttatus* from 10 introduced (UK) and 9 native populations. Within the introduced range we included replicate individuals for four populations (AYR, CER, DBL and HOU), as well as a controlled cross of one of these populations (DBL). For *M. luteus*, we included an inbred line of *M. luteus var. luteus* and one inbred line of *M. luteus var. variegatus* both originally collected in the native range (Chile). The *M. luteus var. luteus* inbred line was represented by two individuals (MLl1 and MLl2, from the 7^th^ and 9^th^ inbred generations). In addition, we analyzed *M. luteus s.l*. from two populations in the introduced range in the UK (COL and EVI), including two individuals originating from the same population (COL): a wild collected plant, and an individual from an experimental cross (Table 2). For *M. x robertsii*, we included samples from four natural populations (GIO, LED, NEN and TOR). For *M. peregrinus* we analyzed samples from three populations (GON, STR, and LED), including three individuals from the type populations for the allopolyploid species (LED). As a control, we also analyzed an experimentally-produced “*M. x robertsii*” hybrid obtained by crossing *M. guttatus* x *M. luteus s.l*. (DBL x COL, maternal x paternal parent; CG-1-1 x CS-4-3 = H003b), as well as an S1 synthetic “*M. peregrinus*” allohexaploid produced by colchicine treatment of the H003b “*M. x robertsii*” line (13-SYN-1-JP1). In addition, we also included three samples from taxa closely related to *M. guttatus* in North America (*M. nasutus, M. micranthus*, and *M. platycalyx*), plus one outgroup (*M, dentilobus*) (Table 2). In terms of sequencing approach, 16 samples (four of each *M. guttatus, M. luteus, M. x robertsii*, and *M. peregrinus*) were derived using sequence capture (Vallejo-Marín et al. 2015), and the remaining through whole-genome sequencing obtained from previously published data (10 samples; Puzey and Vallejo-Marín 2014, 13 samples from the NIH sequence read archive (deposited by JGI; http://www.ncbi.nlm.nih.gov/sra)), or data newly generated here (six samples). In total we analyzed 45 genomic samples (Table 2).

***Genotyping of plastid genomes***—We used the publicly available *M. guttatus* mitochondrial genome (Mower et al., 2012) (JN098455) or the newly assembled *M. luteus var. luteus* chloroplast genome (this study) as references for base calling and genotyping. All reads were aligned to the references using *bowtie2* (Langmead and Salzberg, 2012). Alignments were sorted, duplicates marked and discarded, and read groups added using *Picard* tools (http://broadinstitute.github.io/picard). Following alignment filtration, genotypes were called used GATK UnifiedGenotyper (McKenna et al., 2010; DePristo et al., 2011; Auwera et al., 2013) (Appendix S3; see Supplemental Methods with the online version of this article). Following genotyping, indels and heterozygous sites were excluded.

***Filtering of genomic data for phylogenetic analysis***—The 17 chloroplast-derived regions of the *Mimulus guttatus* mitochondrial genome (NC_018041.1; Mower et al., 2012) were extracted and aligned against the *Mimulus luteus* chloroplast genome using Sequencher v.5.0.1 (http://www.genecodes.com). The *Mimulus luteus* chloroplast sequence was trimmed prior to alignment so that only the LSC, IRB, and SSC were present. Alignments were inspected for sequence similarity. Multiple instances where portions of the mitochondrial chloroplast-derived sequences did not have an ordinal alignment with the majority of the sequence were identified. In these cases, the region that did not align was removed and realigned to the full plastome. These chimeric chloroplast-derived sequences may reflect multiple recombination events, and subsequence recombination hotspots of the mitochondrial genome, or they are representative of multiple chloroplast-genome insertion events in the history of the mitochondrial genome. Once aligned, the coordinates of alignment to the chloroplast genome were recorded. Data corresponding to these positions were eventually removed from the data matrix (see below).

Chloroplast sequences were reconstructed from VCF files representing mapping to the *Mimulus luteus* plastome (see above). For each position of the *M. luteus* plastome, the corresponding base in the mapped data set was recorded. If an indel was identified, that position was ignored. Indels are difficult to distinguish from missing data using this method, so we chose to completely ignore them to reduce potentially conflicting signal. Each reconstructed plastome was split into features (protein-coding genes, tRNAs, and rRNAs) and inter-feature regions, which were used for individual alignments and concatenation (see below). Features were identified based on their alignment to known coordinates of the annotated the *Mimulus luteus* chloroplast genomes. Chloroplast regions identified in mitochondrial genome were excluded in phylogenetic analyses. This process was done using a Perl script (https://github.com/mrmckain/AJB_2015_Mimulus_Allopolyploidy).

Mitochondrial sequences were reconstructed in a similar manner to that of the plastid sequences. In this case, the annotation of the *M. guttatus* mitochondrial genomes was used to recognize features in the reconstructed sequences. Regions mapping to the chloroplast-derived regions of the mitochondrial were ignored. As before, reconstructed features and inter-feature regions were used in alignments and then concatenated (see below).

***Phylogenetic analyses***—Individual features and inter-feature regions were aligned using MAFFT v7.029b (Katoh et al., 2005) on auto settings. Individual alignments were concatenated into a single alignment that was assessed for presence/absence of nucleotides at each site. For the chloroplast alignment, if a gap was identified at a site, that site was removed. As with indels, we chose to do this to alleviate the effects of missing data on the phylogeny estimation. The final, un-gapped alignment was 86,843 bp for all 45 samples. The chloroplast phylogeny was estimated using RAxML v8.0.22 (Stamatakis, 2006) under the GTR+Γ model of evolution with 500 bootstrap replicates.

Mitochondrial sequences were processed in a similar fashion to the plastid sequences. The individual feature and inter-feature regions were aligned using MAFFT and concatenated. The mitochondrial sequence data was missing a larger percentage of the total mitochondrial genome, on average, across the 45 sample set. We noted that the sequence-captured samples were extremely low in their total representation of the mitochondrial genome. Filtering out all gapped sites (as above) resulted in a very short alignment of 4,875 bp. After removal of the sequence-captured samples and the WGS sample CG, which demonstrated a low coverage of the mitochondrial genome, the un-gapped alignment of the remaining 28 accessions totaled 402,717 bp. The mitochondrial phylogeny was estimated as above using the reduced 28 sample set.

We included CS-4-3 (*M. luteus*), CG-1-1 (*M. guttatus*), first generation lab generated triploid hybrid HOO3b (CG-1-1 x CS-4-3), and lab generated allohexaploid SYN (13-SYN-1-JP1) to verify that this approach could determine maternal ancestry in a controlled hybridization event. In addition, we also included two *M. luteus* individuals (MLl1 and MLl2) derived from the same inbred line but genotyped using different methods (WGS and SC, respectively). Alignments and phylogenetic trees are available on Dryad (doi:10.5061/dryad.5q91d).

***Haplotype networks***— We estimated matrilineal haplotype networks for both chloroplast and mitochondria using statistical parsimony (TCS) implemented in PopArt (http://popart.otago.ac.nz). For this analysis we used a panel of 929 SNPs for the chloroplast, and 1,454 for the mitochondria. Only variable sites were included in the SNP matrix. Indels and sites filtered as described in above section (*Filtering of genomic data for phylogenetic analysis*) were excluded. For chloroplast, we included both whole-genome sequence (WGS) and sequence capture (SC) samples, and for mitochondria we excluded the SC samples as above. For both data sets we excluded sites with more than 5% missing data. This allowed us to analyze 493 sites of which 161 were parsimony-informative for chloroplast, and 784 sites of which 300 were parsimony-informative for the mitochondria.

***Principal component analysis***—We also conducted a principal component analysis (PCA) using the SNP data of the chloroplast and mitochondrial genomes using the *glPca* function in the *R* package *adegenet* v. 2.0 (Jombart and Ahmed, 2011). For this analysis we removed genomic regions with low genotyping success, and included only those SNP sites that were successfully genotyped in 90% (41/45) or more individuals. This filtering step removed poorly genotyped genomic regions, and reduced the total number of SNPs from 1454 to 434 for the mitochondrial genome and from 929 to 694 in the chloroplast. For the chloroplast analysis, we also removed a single WGS individual which had poor genotyping success across the genome even after filtering (CG). As expected the amount of missing data across genotyped SNP loci was higher for the SC than for the WGS data set after filtering. For the mitochondria data set, the amount of missing data per site per individuals was 0.0013 for WGS and 0.1080 for SC. For the chloroplast, the same threshold yielded an average amount of missing data per site per individuals of 0.0054 for WGS and 0.0176 for SC.

***Mitochondrial genotyping in natural hybrids***—We used mitochondrial genome sequences to identify potential markers that could help us distinguish *M. guttatus* and *M. luteu*s in natural hybrid populations. We sought loci segregating for alternative alleles in *M. luteus* (MLl1 and CS) and *M*. guttatus (CG and AYR1) in the sequenced panel. Our goal was to identify fragment length polymorphic sites (indels) that could be scored using microcapillary fragment analysis. To identify segregating indels, we sorted BAM alignments in IGV genome browser, and manually screened for indels that were alternatively fixed between *M. luteus* and *M. guttatus*. We identified three indels (3-4 bp) that distinguished *M. guttatus* and *M. luteus*. Genotyping individuals at these indels produced a distinct haplotype with which we could potentially identify the maternal (mitochondrial) parent of hybrid individuals. To facilitate large-scale genotyping, we designed primers flanking each of the indel regions using Primer3Plus (Untergasser et al., 2012), and confirmed their amplification, and sequence in a separate test panel of parental and hybrid individuals using Sanger sequencing. We checked the compatibility of the three primer pairs for pooling them in a single multiplex reaction using Multiplex Manager v 1.2 (Holleley and Geerts, 2009). We then used fluorescently-labelled forward primers (6-FAM, Eurofins Genomics, Germany), and unlabeled reverse primers to generate PCR products in a single multiplex reaction (Type-It, QIAGEN, UK). PCR cycles consisted of a denaturing step of 5 min at 95 ° C, followed by 30 cycles of 95 ° C for 30 s, 55 ° C for 180 s, and 72 ° C for 30 s, and a final elongation step of 30 min at 60 ° C. Fragment length of the PCR products was measured in an ABI3730 sequencer using LIZ500 as a size standard (DNA Sequencing and Services, Dundee, UK). Primer sequences for these mitochondrial markers are given in Appendix S4 (see Supplemental Data with the online version of this article). We genotyped 163 individuals of *M. guttatus* (3 populations), *M. luteus s.l*. (4 populations), *M. x robertsii* (13 populations), and *M. peregrinus* (2 populations), as well as a synthetic allohexaploid line product of an *M. guttatus* (DBL) x *M. luteus* (COL) cross treated with colchicine and selfed for one generation (SYN; Table 2). Between 1 and 15 individuals were genotyped per taxon per population (6.8 ± 3.5, mean ± SD) (Table 3).

**Table 3.**
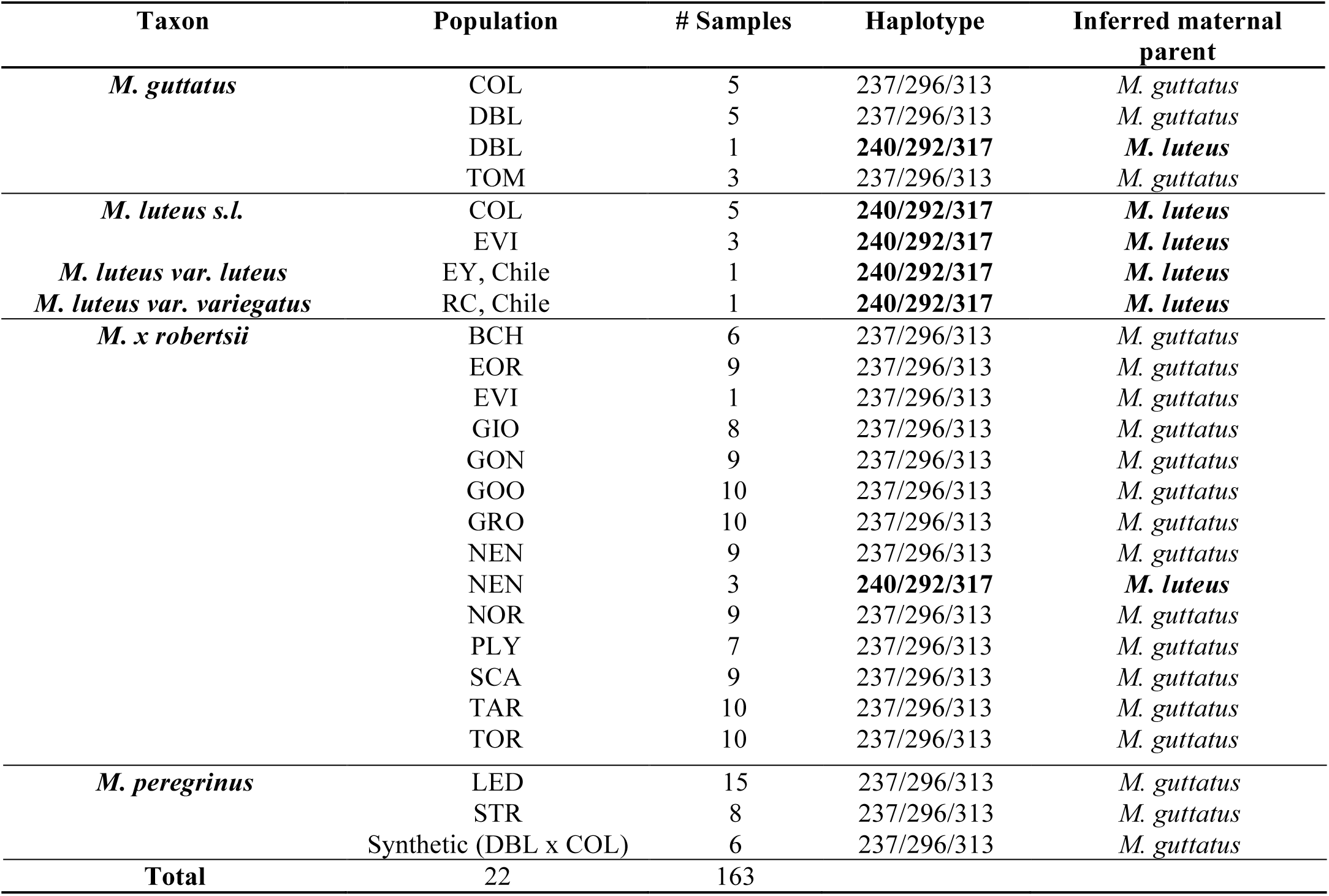
Mitochondrial haplotypes of 163 individuals of *M. guttatus, M. luteus*, and their hybrid derivatives, the triploid *M. x robertsii* and allopolyploid *M. peregrinus*. Haplotypes are given as the size in base pairs of three mitochondrial loci with fragment-length polymorphism (indels). Between 1 and 15 individuals of each of 21 populations, and one artificially created allopolyploid, were genotyped at all three loci. The synthetic allopolyploid was the product of a *M. guttatus* (DBL) x *M. luteus s.l*. (COL) cross, with *M. guttatus* as the maternal parent. A single
individual phenotyped as *M. guttatus* (DBL), but with a “*M. luteus*” maternal haplotype is presented separately.

## RESULTS

***Seed set and germination of artificial hybrids***—Reciprocal crosses between *M. guttatus* and *M. luteus* resulted in a relatively high number of seeds per fruit in both directions. The average number of seeds per fruit was 723 ± 73 (mean ± SE) for the MG x ML cross, and 635 ± 100 for ML x MG. This difference was not statistically significant (Tukey test, *P* = 0.72; Fig. 1A). Within species crossing yielded 1,237 ± 95, and 467 ± 57 seeds per fruit for *M. guttatus* and *M. luteus*, respectively (Fig. 1A). In comparison, we found a marked effect of the identity of the maternal parent on seed germination. While approximately half of the seeds of the MG x ML cross germinated (0.525 ± 0.062), only 4% of the ML x MG were able to do so (0.042 ± 0.018) (Fig. 1B). Both parental taxa had high germination proportions (0.835 ± 0.037 and 0.825 ± 0.020, for MG x MG, and ML x ML, respectively), although *M. guttatus* seeds achieved maximum germination at a much faster rate (Fig. 1B).

***Floral and vegetative phenotype of F1 hybrids***—The proportion of hybrid plants that flowered was significantly different depending on crossing direction. The MG x ML cross had a higher proportion of flowering individuals (0.983 ± 0.017, *n* = 60) than the reciprocal cross (0.706 ± 0.114, *n* = 17) (Tukey test *z* = 2.809; *P* = 0.021). The proportion of flowering plants for the MG x ML cross also exceeded both parental crosses (0.633 ± 0.044, *n* = 120; and 0.741 ± 0.040, *n* = 120; for MG x MG (*z* = 3.442, *P* < 0.01) and ML x ML (*z* = 2.979; *P* < 0.05), respectively). Moreover, MG x ML hybrids began flowering on average nine days earlier than the reciprocal cross (54.91 ± 0.96 days, 63.25 ± 4.08). Flowering occurred slightly earlier for *M. luteus* (53.08 ± 0.98) than for *M. guttatus* (57.77 ± 1.14).

Among plants that flowered, the MG x ML hybrid achieved the largest stature of all cross types (267 ± 9mm, *vs*. 139.1 ± 12.3, 188.7 ± 5.0, and 138.9 ± 5.4; for ML x MG (*z* = 7.491, *P* < 0.001), MG x MG (*z* = 8.431, *P* < 0.001), and ML x ML (*z* = 14.011, *P* < 0.001), respectively; Fig. 2). Flowers and bracts of both types of hybrids were intermediate or larger than the parental values (Table 4, Appendix S5-S6 (see Supplemental Data with the online version of this article)). Both MG x ML and ML x MG hybrids had similar-sized flowers, although hybrids with *M. guttatus* as the maternal parent had longer pedicels and larger bracts (Table 4). Parental taxa differed significantly in nectar production (ML x ML produced more nectar than MG x MG; Tukey test *z =* 4.998; *P* < 0.001), but while MG x ML hybrids produced significantly more nectar than MG x MG (*z* = 6.138; *P* < 0.001), ML x MG hybrids produced larger amounts, but not significantly so (*z* = 2.299; *P* = 0.092), than the *M. guttatus* parent (Fig. 2).

**Figure 2.**
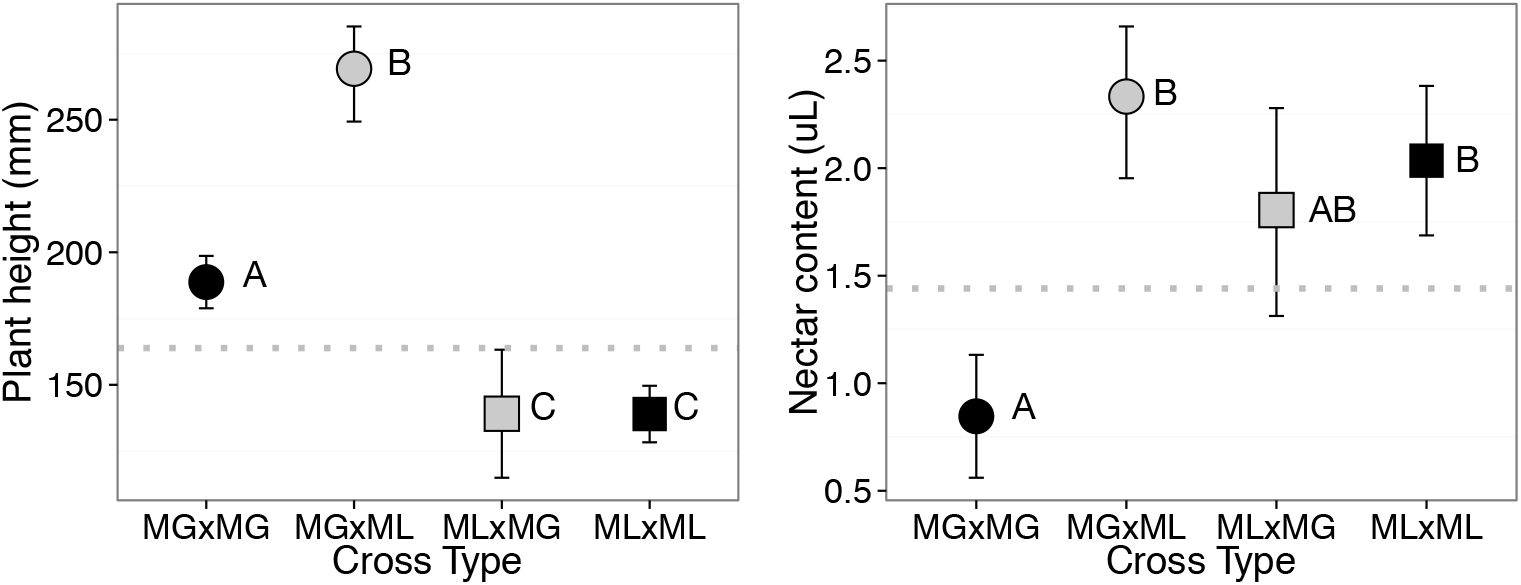
Plant height at first flower **(A)**, and nectar content per flower (**B**) of *Mimulus* crosses within- (MG x MG and ML x ML) and between-species (MG x ML and ML x MG). Sample size (number of individuals) for plant height: *n* = 72, 54, 11, 78; sample size (number of flowers): *n* = 67, 73, 14, 73. Dotted line shows the mid-parent value. Bars are 95% confidence intervals.

**Table 4.**
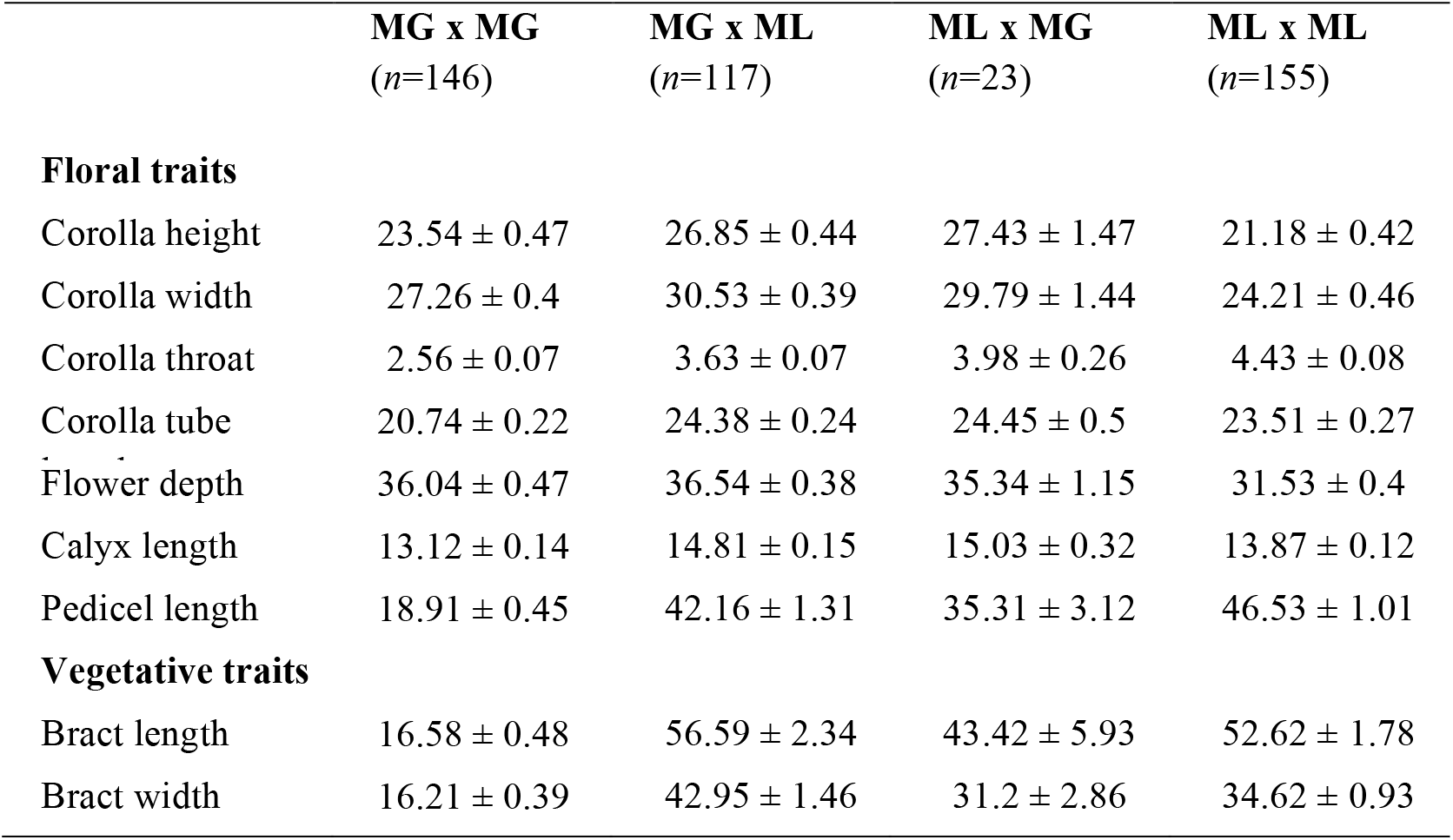
Phenotypic measurements of *M. guttatus* (MG x MG), *M. luteus* (ML x ML), and their reciprocal hybrids (MG x ML, and ML x MG). Crosses are shown as maternal x paternal parent. Mean +/- standard error (SE). All values in millimeters.

***Analysis of Mimulus chloroplast***—The complete *Mimulus luteus* chloroplast genome was assembled, providing the first published sequence for both the genus *Mimulus* and the family Phrymaceae. The plastome follows the typical quadripartite structure found in most of the photosynthetic angiosperm lineages. The total length of the chloroplast genome is 153,150 bp and is made up of a 84,293 bp LSC, a 17,851 bp SSC, and two 25,503 bp IR regions (Fig. 3). The complete chloroplast sequence is available on Genbank (KU705476). The plastome contains 80 unique protein-coding genes (including *ycf1*, *ycf15*, and *ycf2*), 30 unique tRNAs, and 4 unique rRNAs. No major gene losses or rearrangements were identified relative other published plastomes from Lamiales (Qian et al., 2013; Zhang et al., 2013; Nazereno, Carlsen, and Lohmann 2015.

**Figure 3.**
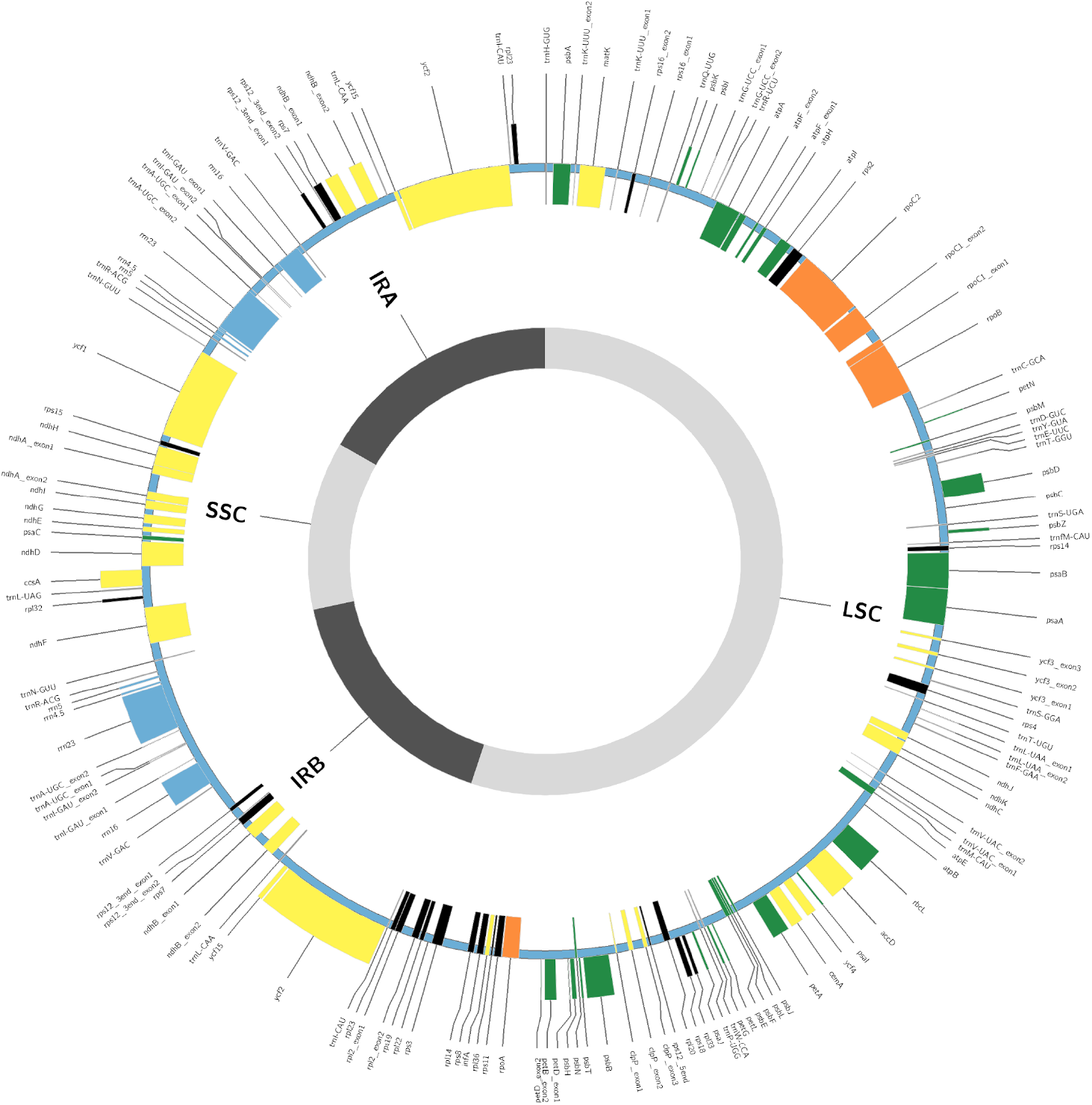
Circular plot of *Mimulus luteus* var. *luteus* chloroplast genome. The plastome is 153,150 base pairs with a 84,293 base pair large single copy (LSC) region, a 17,851 base pair small single copy (SSC) region, and two 25,503 base pair inverted repeat regions (IRB and IRA). Photosystem genes are in green, RNA polymerase subunits are in orange, ribosomal genes are in black, other protein encoding genes are in yellow, rRNAs are in blue, and tRNAs are in black.

***Genomic analysis of hybrid parentage based on cytoplasmic genomes***— On average, sequencing coverage of the cytoplasmic genomes was very high, although also very skewed. For WGS datasets, per individual average read coverage of the mitochondrial and chloroplast genome ranged from 48-940x and 347-5556x with global means of 318 and 2303, respectively. As expected, the sequence capture datasets had high read coverage but totals were considerably lower than the WGS datasets (Appendix S2). Per individual average read coverage for SC datasets mapped against the mitochondrial and chloroplasts genomes from 6-13x and 55-112x with means of 9x and 85x, respectively. The chloroplast has considerably greater read depth than the mitochondria. The fraction of genotyped bases varied considerably between the WGS and SC dataset (Appendix S7; see Supplemental Data with the online version of this article). On average, 68% and 80% of chloroplast sites were genotyped in the SC and WGS datasets, respectively (fraction of sites genotyped calculated from entire mitochondrial and chloroplast datasets before filtering). The mitochondria exhibited larger span between the average percentage of called bases between the SC and WGS datasets with 57% and 85% called.

The chloroplast tree had relatively high resolution and strong support at the taxon level (i.e., *M. guttatus vs. M. luteus*), but was less useful to resolve relationships between populations within species (Fig. 4), suggesting recent divergence of populations. A single monophyletic *M. luteus* clade (minus MLv) is well supported in the chloroplast tree. The mitochondrial dataset does not support *M. luteus* monophyly. Within the chloroplast tree, *M. peregrinus* (allopolyploid) forms two distinct groups and are more closely related to geographically proximate sterile triploid hybrids. *M. x robertsii* sample (LED1), and *M. peregrinus*, LED2, LED3, and LED4, are closely related and all collected form Leadhills, Scotland. *M. peregrinus* sample GON, also falls within the above group and is from nearby Glengonnar, Scotland. Based on the chloroplast data, the other naturally *M. peregrinus* sample included in this analysis, STR, also falls within a clade with geographically proximate *M. x robertsii* (TOR). The relationship of *M. x robertsii* and *M. peregrinus* supports the finding of Vallejo-Marín *et al*. (2015) using a different aspect of the genomic data, i.e., the nuclear component of the genome. Naturally occurring allopolyploids (LED2, LED3 LED4, GON, and STR) as well as naturally occurring hybrids (LED1, TOR, NEN, GIO) are more closely related to *M. guttatus* than they are to *M. luteus* based on chloroplast analysis. The mitochondrial tree topology indicates that naturally occurring allopolyploid LED is nested within a *M. guttatus* clade. The chloroplast tree places MLl1 and MLl2 (samples derived from the same inbred lines but genotyped using different methods) sister to each other. SYN and HOO, triploid hybrid and allohexaploid samples, respectively, derived from an artificial cross between CG x CS, fall within the same clade in both the chloroplast and mitochondrial trees. The maternal parent of these lines (CG) is not sister to the SYN and HOO samples as expected. This is most likely due to homoplasy affecting tree reconstruction due to relatively low sequence divergence between accessions.

**Figure 4.**
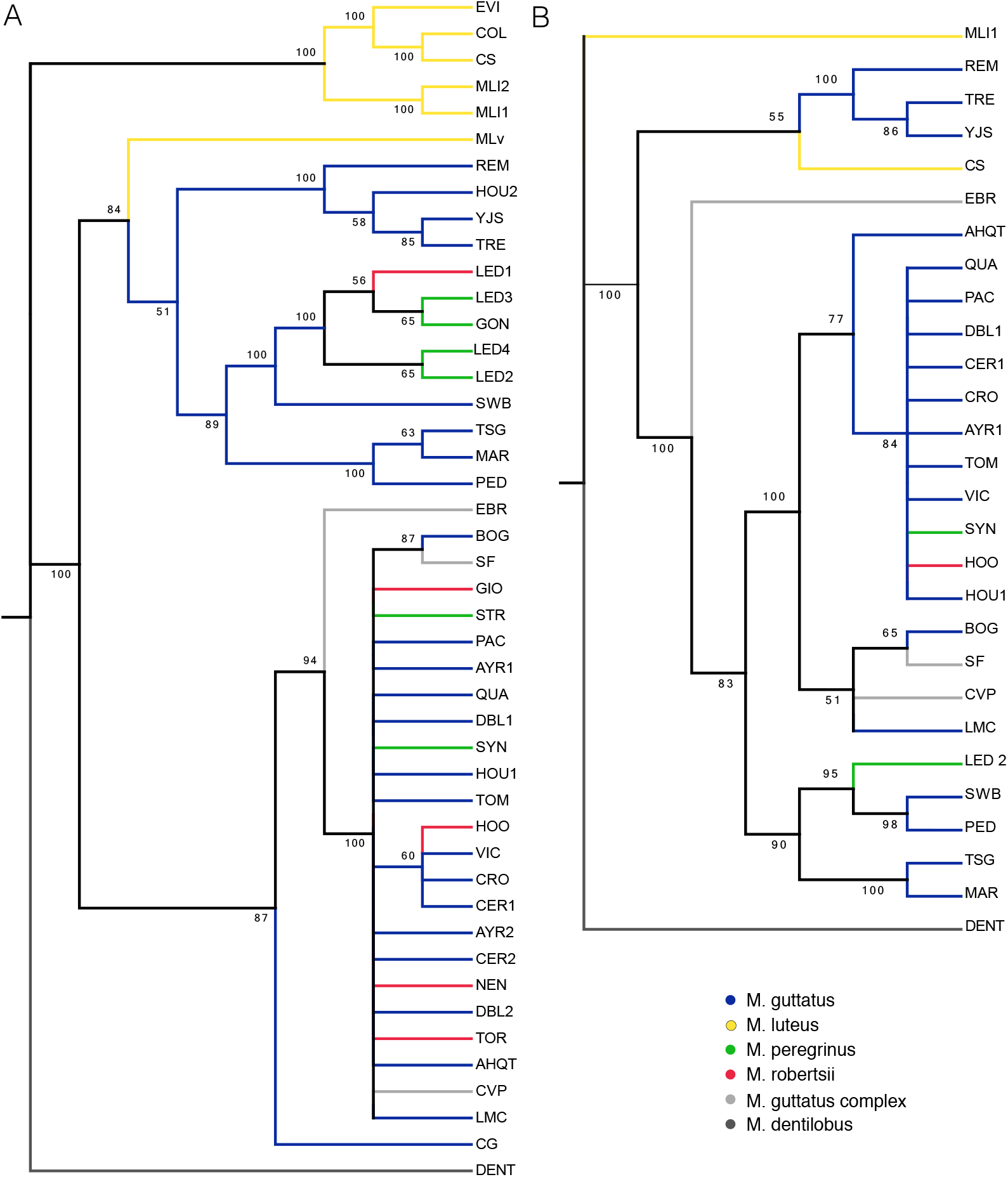
Maximum likelihood tree of *Mimulus* species and populations based on alignments of 86,843 base pairs from chloroplast genomes (A) and 402,717 base pairs from mitochondrial genomes (B). Bootstrap values are indicated. Nodes with less than 50% bootstrap support were collapsed. Samples genotyped using sequence-capture (N=16) are excluded from the mitochondrial analysis due to low genotyping success. Trees are rooted with *M. dentilobus*, Color-coding is based on species identity.

The haplotype network of the chloroplast showed that both *M. x robertsii* and *M. peregrinus* samples have identical or nearly identical haplotypes to *M. guttatus* (Fig. 5A). Samples of *M. peregrinus* from Leadhills (LED2, LED3, LED4) and nearby Glengonnar (GON) share a haplotype with *M. x robertsii* from Leadhills (LED1). Samples of *M. peregrinus* from Orkney (STR) belong to a closely related haplotype that also includes *M. x robertsii* from Orkney (TOR), northern England (rob3), and the Outer Hebrides (rob4). This haplotype also includes most of the *M. guttatus* samples from the British Isles, the synthetic triploid and hexaploid (HOO and SYN), and one native population (AHQT). Samples of *M. luteus s.l*. from the British Isles (CS and COL, Coldstream; EVI, Evie) fall in a distinct group of haplotypes separated from other *Mimulus* by several mutational steps, and which also includes the two replicates from the *M. luteus var. luteus* inbred line (MLl1 and MLl2; Chile). As in the phylogenetic reconstruction, the sample from a Chilean *M. luteus var. variegatus* (MLv) is situated near *M. guttatus* haplotypes. The mitochondrial network, which excludes all SC samples due to low genotyping success, shows the single individual of wild *M. peregrinus* (LED2) as part of a haplotype that includes most of the British *M. guttatus* as well as the synthetic triploid and hexaploid accessions (HOO and SYN). Both *M. luteus var. luteus* (MLl1) and *M. luteus s.l*. (CS) are separated from all other *Mimulus* by a large number of mutational steps (Fig. 5B).

**Figure 5.**
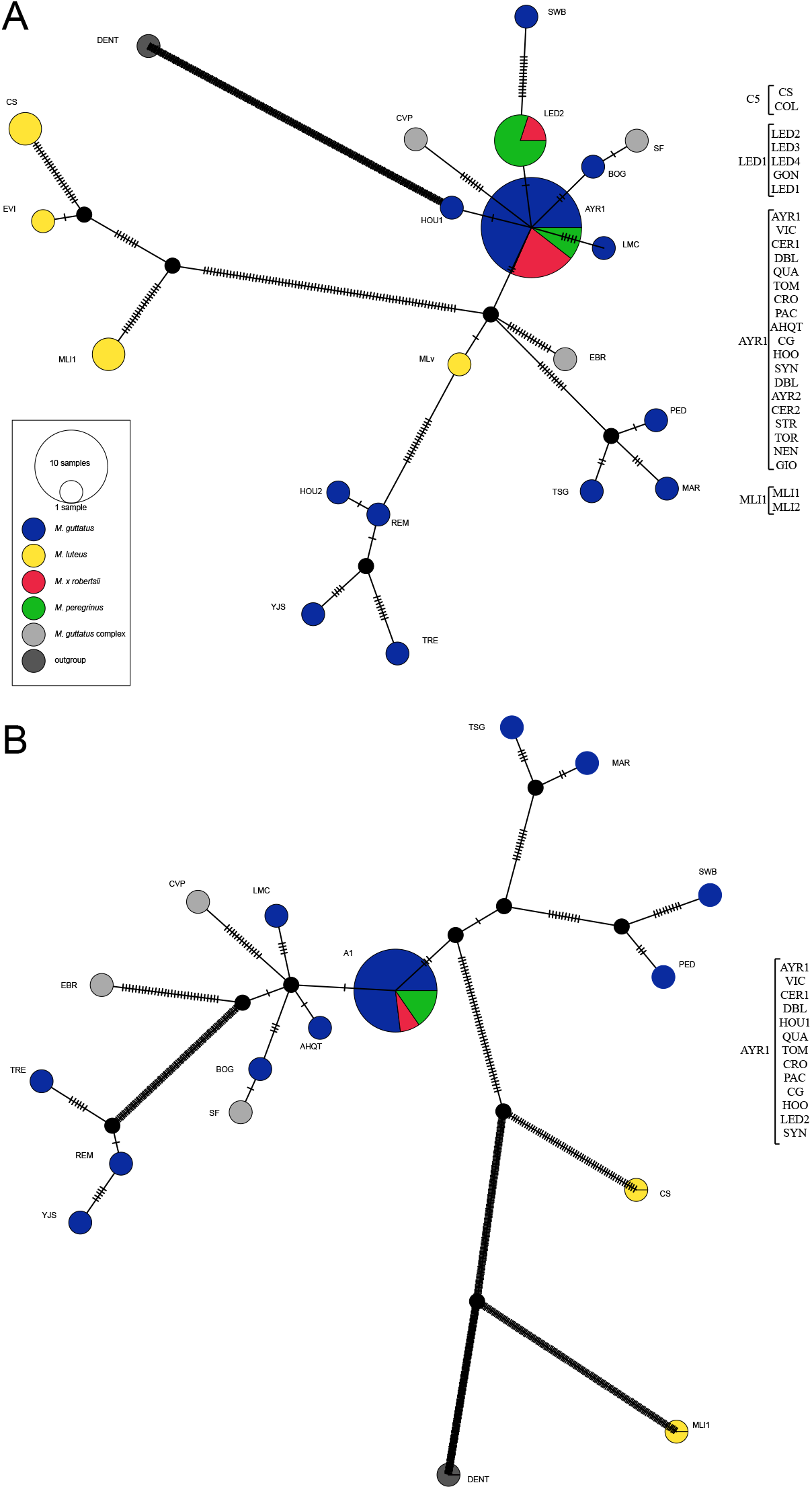
Matrilineal haplotype network (statistical parsimony; TCS) of (A) chloroplast and (B) mitochondrial SNPs. The chloroplast analysis is based on 493 sites, and the mitochondrial on 784 sites. Sites with more than 5% missing data were excluded. Hash marks on lines are mutational steps. The haplotype circles are labeled with one individual and all other individuals with that haplotype are listed to the right. The mitochondrial dataset excludes 16 samples obtain through sequence capture due to low genotyping success (see Results). Names for individual samples are only shown here for *M. x robertsii* and *M. peregrinus*.

The PCA of both chloroplast and mitochondria data sets also showed that *M. x robertsii* and *M. peregrinus* cluster together with *M. guttatus* samples (Fig. 6). In the chloroplast PCA, the two known origins of *M. peregrinus* (Leadhills and Orkney) fall in separate clusters with *M. guttatus*, each of them associated with local *M. x robertsii* samples (Fig 6A, Appendix S8; see Supplemental Data with the online version of this article). In the mitochondrial PCA, there is not enough resolution to differentiate these two origins, and all *M. peregrinus* and *M. x robertsii* samples fall in the same cluster, along with other native and introduced *M. guttatus* (Fig. 6B, Appendix S9; see Supplemental Data with the online version of this article). As above, the *M. luteus var. variegatus* sample from Chile (MLv) is closer to *M. guttatus* samples than to other *M. luteus*.

**Figure 6.**
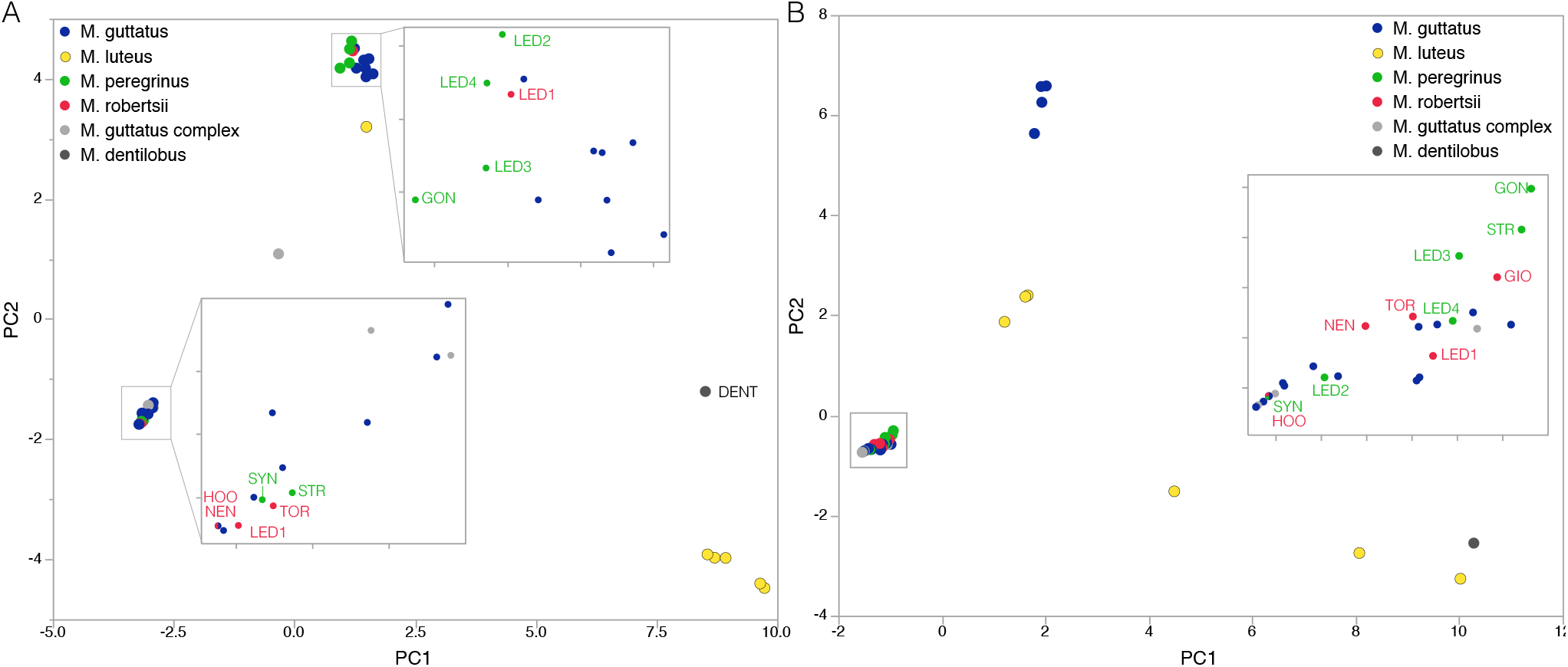
Principal component analysis of (A) 929 SNPs from the chloroplast and (B) 1454 SNPs from the mitochondria. Only sites successfully genotyped in 41/45 samples were included in this analysis. The chloroplast dataset excludes one individual (CG) with low genotyping success for the analyzed SNPs. The mitochondrial dataset excludes 16 samples obtain through sequence capture due to low genotyping success (see Results). A fully annotated version of this Figure is available as Appendix S8 and Appendix S9.

***Natural frequency of* M. guttatus *as maternal parent***—We successfully genotyped 163 *Mimulus spp*. individuals from 21 populations and one synthetic line at three mitochondrial loci (Table 3). We recovered only two haplotypes: *237/296/313* and *240/292/317* (fragment size in base pairs for markers mit-259, mit-262, and mit-357, respectively). For *M. luteus*, we genotyped 10 individuals, all of which had haplotype *240/292/317*. For *M. guttatus*, we genotyped 14 individuals from three populations of which all but one had haplotype *237/296/313*. The exceptional individual belonged to population DBL (Table 3). We genotyped 110 individuals of *M. x robertsii* from 12 populations across the U.K. From these, 107 had haplotype *237/296/313* (*“M. guttatus* haplotype”), while three individuals from population NEN had the other haplotype. Finally, the 23 individuals of *M. peregrinus* sampled in populations STR and LED also had the haplotype *237/296/313*. The synthetic allohexaploid line created with *M. guttatus* as the maternal parent, had haplotype *237/296/313* as expected.

## DISCUSSION

Our study combines experimental crosses, genomic analyses of chloroplast and mitochondrial genomes, and surveys of natural hybrid populations to demonstrate how a strongly asymmetric post-zygotic hybridization barrier has shaped the origin of the recently formed allopolyploid species *Mimulus peregrinus*, and its triploid ancestor *M. x robertsii* (= *M. guttatus* x *M. luteus*). Thus our study confirms early crossing experiments (Roberts, 1964; Parker, 1975), and adds to a growing list of exceptions to the “rule” of poorly-performing paternal-excess crosses (Table 1; Appendix S1; see Supplemental Data with the online version of this article). Specifically, we showed that viable hybrids are considerably more likely to be produced when diploid *M. guttatus* is used as the maternal parent and tetraploid *M. luteus* as the paternal parent (MG x ML), than in the opposite direction (ML x MG). We showed that MG x ML hybrids are larger in size than the parental taxa and the reciprocal hybrid ML x MG. The MG x ML hybrids also flowered earlier and produced more nectar than their *M. guttatus* parents. Motivated by this observed asymmetry of hybridization, we used previously available genomic sequences, and developed novel genomic resources, including the first assembled chloroplast in the genus *Mimulus* and in the family Phrymaceae, to show that most individuals in natural populations of the triploid hybrid *M. x robertsii* have *M. guttatus* as their maternal parent. As expected, *M. peregrinus*, which arose by genome duplication from *M. x robertsii*, also has *M. guttatus* as its maternal parent. Beyond revealing the asymmetric origin of *M. x robertsii* and *M. peregrinus*, our study shows the general potential of exploiting cytoplasmic genome data from both targeted and whole-genome sequence projects (Straub et al., 2012; Dodsworth, 2015), to investigate the evolution and speciation of polyploid and hybrid taxa.

***Pattern of hybridization asymmetry: seed viability and germination***—Cytonuclear incompatibilities and differences in ploidy level, or a combination of both, can result in post-zygotic asymmetric hybridization (Tiffin, Olson, and Moyle, 2001; Köhler, Scheid, and Erilova, 2010; Scott, Tratt, and Bolbol, 2013). We currently cannot distinguish between these two classes of mechanisms to explain the observed pattern of asymmetric hybridization leading to *M. x robertsii* and *M. peregrinus*. However, analysis of the morphology of hybrid seeds obtained in reciprocal crosses may offer some insight into the mechanisms of such asymmetry. For example, intra-specific hybridization between diploid and tetraploid *Arabidopsis thaliana* has shown that crosses with maternal excess (4x – 2x) yield smaller seed sizes than the reciprocal paternal excess cross (2x – 4x) (Scott, Tratt, and Bolbol, 2013). The seed size differences are thought to result from misregulation of imprinted genes that control growth and cell division in the endosperm (Köhler and Kradolfer, 2011; Schatlowski and Köhler, 2012; Lafon-Placette and Köhler, 2015). Paternal excess may cause endosperm over-proliferation and failed endosperm cellularization (resulting in large seeds), while maternal excess results in endosperm proliferation and small seeds (Haig and Westoby, 1991; Scott et al., 1998; Bushell, Spielman, and Scott, 2003; Scott, Tratt, and Bolbol, 2013).

As a first step toward understanding the mechanism of reproductive asymmetry in *Mimulus*, we compared seed size in the 16 controlled crosses used in the phenotypic experiment. These crosses included four accessions of each parental taxa and the reciprocal hybrid crosses (Appendix S2; see Supplemental Data with the online version of this article). We estimated seed size as seed area obtained from digital images of 300-600 seeds per cross type (Appendix S10; see Supplemental Data with the online version of this article). Our results indicate that although both types of hybrids have smaller seeds than the parental taxa, the ML x MG cross produces larger (if mostly inviable) seeds than the reciprocal (and more viable) MG x ML cross (Appendix S10; see Supplemental Data with the online version of this article). This suggests that interploidy crosses in *Mimulus* do not mimic the seed phenotypes of interploidy crosses in *Arabidopsis* (Scott, Tratt, and Bolbol, 2013). Further analyses of the morphology and histology of interploidy reciprocal crosses of *M. guttatus* x *M. luteus* are needed to elucidate the mechanistic and genetic basis of post-zygotic hybridization asymmetry in this group.

***Pattern of hybridization asymmetry: adult plant phenotype***—Most previous experimental studies comparing the viability of asymmetrically produced hybrids have focused on the immediate consequences of intrinsic post-zygotic incompatibilities at the seed and seedling stage (seed number and seed germination). However, as we have shown, surviving hybrids may also differ in their adult phenotypes depending on cross direction. Although our sample size is small, our results show that paternal-excess *Mimulus* triploids possessed characteristics potentially associated with high extrinsic fitness, including larger plant size. Paternal excess hybrids also flowered earlier, and produced more nectar than maternal-excess triploids, but the direct effects of these traits on hybrid fitness are expected to be negligible given that *M. x robertsii* is sexually sterile (Roberts, 1964; Stace, 2010), and this sterility does not differ in synthetic hybrids created using reciprocal crosses (Roberts 1964, p. 73). A major gap in our knowledge is whether phenotypic differences among reciprocal hybrids observed in greenhouse conditions, translate to performance differences in the field, and further studies on the ecology and fitness of reciprocal hybrids and their parents are needed. It would also be important to compare the morphology of triploid *M. x robertsii* and its allohexaploid derivative *M. peregrinus*, in order to assess the effects of genome duplication on the phenotype. Preliminary field observations suggest that allopolyploids are larger in size for some morphological traits such as flower size (Vallejo-Marín, pers. obs.). The combination of *Mimulus* genomic and transgenic resources (Wu et al., 2008; Hellsten et al., 2013; Yuan et al., 2013; Twyford et al., 2015) with a rich ecological knowledge base, makes the naturalized hybrids and neopolyploids of the British Isles an exciting model for connecting molecular mechanisms of interploidy asymmetry to observed patterns of speciation in the wild.

***Genetic ancestry of* M. x robertsii *and* M. peregrinus**—Regardless of mechanism, our study seems to be an exception to the general pattern in which viable inter-ploidy hybrids are more likely to be produced when the maternal parent is of higher ploidy level than the paternal one (maternal excess; Stebbins, 1957; Ramsey and Schemske, 1998). Both chloroplast and mitochondrial genomic analyses clearly show that the triploid and hexaploid individuals cluster more closely with *M. guttatus* than with any native or introduced *M. luteus* (Figs. 4, 5, 6). The genomic data also shows that even whole-genome sequences are unable to completely separate *M. guttatus* and *M. luteus*, which may reflect incomplete lineage sorting between these relatively closely related taxa (Madison and Knowles, 2006) Moreover, the origin of *M. luteus* itself is unknown, but nuclear genomic data suggests that this taxon has a hybrid origin (Vallejo-Marín et al., 2015), perhaps with a *M. guttatus*-like ancestor (Mukherjee and Vickery, 1962). If the putative allopolyploid *M. luteus* has had multiple origins, then the simple expectation of reciprocal monophyly for the plastid genomes of *M. luteus* and *M. guttatus* may not be met. Solving the mystery of the origin of *M. luteus* and its relationship with *M. guttatus* and related taxa will require detailed analyses of natural populations throughout the native South American range.

Our survey of hybrid *Mimulus* populations across the British Isles using three mitochondrial loci also supports *M. guttatus* as the maternal parent of *M. x robertsii* and *M. peregrinus*. Of 114 triploid *M. x robertsii* from 13 populations, and 27 individuals from three populations of the hexaploid *M. peregrinus* that were genotyped at these three mitochondrial loci, all but three *M. x robertsii*, individuals from a single population had maternal haplotypes that could be confidently assigned to *M. guttatus*. The three individuals of *M. x robertsii* (all from population NEN) that had *M. luteus* mitochondrial haplotypes (Table 3) pose an exception to the general pattern detected here. An *M. luteus*-like maternal haplotype in hybrids could be explained in a number of ways. First, it could simply be due to contamination during genotyping. However, we re-extracted and re-genotyped this samples and confirmed the mitochondrial haplotype. Second, the three samples could have been mistakenly identified as *M. x robertsii* but instead belong to *M. luteus*. We think this is also unlikely as plants were morphologically identified on site, and we have also conducted surveys of this population (NEN) in 2010, 2011, and 2013, and *M. luteus* has never been observed in this or nearby populations. Third, the haplotypes in these three hybrids could arise via introgression from *M. luteus*. Introgression across ploidy barriers has been shown for nuclear markers in other species (Chapman and Abbott, 2010), and could possibly occur at the mitochondrial level as well. Although, we cannot rule out this possibility, the lack of local *M. luteus* (the nearest known *M. luteus* natural population is ˜100km north), and principally the fact that hybrids are not known to set seed (i.e., are almost completely sterile; Roberts, 1964; Vallejo-Marín, 2012), makes the introgression scenario unlikely. Fourth, the “*M. luteus*” haplotype in hybrids could be explained by incomplete lineage sorting of ancestral polymorphisms between *M. guttatus* and *M. luteus* (Twyford and Ennos, 2012). We cannot currently rule out ancestral polymorphism, and this remains a distinct possibility given the observations that a single individual of *M. guttatus* (DBL) bears a mitochondrial haplotype otherwise characteristic of *M. luteus* (Table 2). Similarly the mitochondrial haplotype for the British individual of *M. luteus s.l*. (CS) nested within the *M. guttatus* clade also suggests incomplete lineage sorting. Given the limited sampling of parental genotypes, incomplete lineage sorting remains a very likely possibility. Fifth, it is possible that homoplasy resulting from the sequences converging on a similar haplotype (especially if the sequences are not that distinct) could explain this pattern. Finally, a simpler explanation may be that although the majority of hybrids are produced when *M. guttatus* is the maternal parent, occasionally, the opposite cross can yield viable offspring. Indeed, our experimental results show that viable hybrids in the ML x MG direction are produced albeit with low probability (Figure 1).

**Figure 1.**
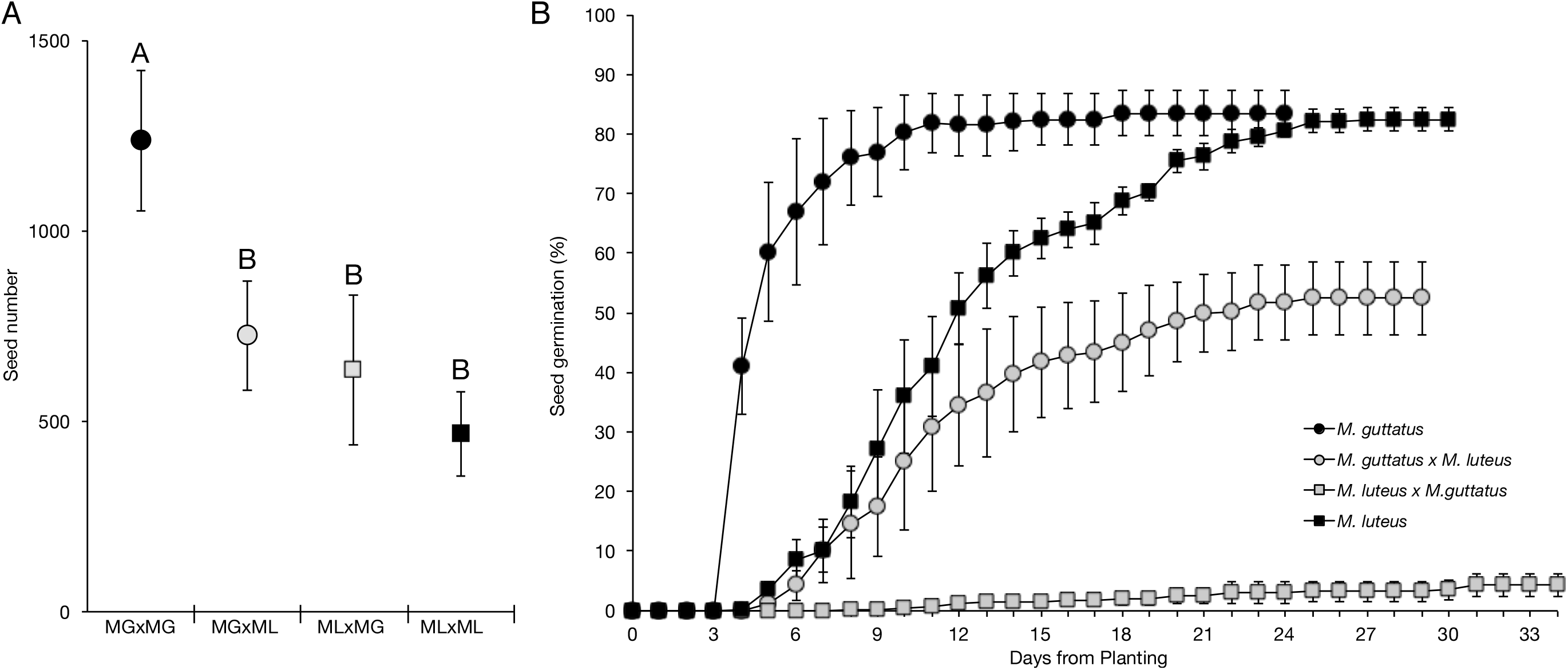
**(A)** Mean seed number and 95% confidence intervals (CI) obtained in crosses within species (MG x MG, and ML x ML), and between species (MG x ML and ML x MG). Sample sizes: 9, 11, 10, 10 fruits. Different letters indicate significantly different means as assessed with a pairwise Tukey test (*z* > 2.98, *P* < 0.05). **(B)** Mean germination of *M. guttatus* x *M. guttatus* (black circle), *M. luteus* x *M. luteus* (black square), *M. guttatus* x *M. luteus* (gray circle), *M. luteus* x *M. guttatus* (gray square). Crosses are indicated as maternal x paternal parent. Four lines for each cross type were analyzed, planting 100 seeds per line (1600 seeds total). Each data point represents the average (+/- SE) proportion of germinated seeds across the four lines for a given cross type across the germination period.

***Skimming of cytoplasmic genomes*** — Acquisition of complete organellar genomes is increasingly feasible with the advent of “genome skimming” techniques associated with next-generation sequencing projects. In fact, mitochondrial sequences usually represent 1-5% of the reads obtained in such projects (Steele, et al., 2012; Smith, 2015) and are often discarded as contamination during nuclear genomic analyses. Our study showed that data generated by whole-genome sequencing contain enough cytoplasmic ‘contamination’ to yield between 1,515 and 208-fold mean coverage of the chloroplast and mitochondria, respectively. Even the approach of sequence capture, in which specific (nuclear) regions were targeted (Vallejo-Marín et al., 2015) contained sufficient reads to build large contigs of both chloroplast and mitochondrial genes (86,843 bp and 402,717 bp, respectively). As expected, however, the sequence capture data did not allow us to genotype all the same sites in all individuals. The limited coverage of specific sites was particularly acute for the mitochondrial genome, which reduced its utility in some of the analyses conducted here. Our results suggest that using off-targeted reads from sequence capture approaches may be sufficient to generate moderate numbers of genetic markers (e.g., Fig. 6), but this approach is of limited utility when coverage of large numbers of sites across multiple individuals is required (e.g., Figs. 4, 5).

***The* Mimulus *chloroplast genome***—Shallow whole-genome sequencing (also referred to as genome skimming) allowed us to reconstruct the chloroplast genome of *M. luteus*. The structure and gene content of this genome follows what has been seen for other taxa in Lamiales (Nazareno et al., 2015), including that of *Utricularia gibba* (Ibarra-Laclette et al., 2013), a species known for a diminutive nuclear genome. The relatively constant architecture and gene content identified across multiple lineages of the order Lamiales suggests that there may be selection for maintenance of gene content and structure relative to other eudicots lineages like saguaro (Sanderson et al., 2015) and legumes (Saski et al., 2005). A more comprehensive taxon sampling of chloroplast genomes is needed throughout the Lamiales and all angiosperm to truly understand the dynamics and evolution of this vital organelle.

The use of genome skimming has repeatedly proven to be a useful technique for acquiring either complete gene sets (e.g. Washburn et al., 2015) or full plastomes (e.g. Bock et al., 2013). Here we demonstrate the utility of a single plastome and mitochondrial genome as anchors to identify organellar genome sequences rapidly from genome skimming data. Though full plastid and mitochondrial genome sequences were not recovered, a large proportion of each genome was and those regions were identified to annotated features and alignable providing resolution of relationships at the population level. Future work in the group will include *de novo* assembly of whole chloroplast genomes across the genus providing valuable resources for population-level studies of hybridization and polyploidy.

***Conclusions***—The value of genome skimming for general phylogenetic analyses is well recognized (e.g. Hahn, Bachmann, and Chevreux, 2013). We demonstrate that the analysis of the entire mitochondrial and chloroplast genomes can yield information to infer the maternal ancestry of a hybrid and neo-allopolyploid derived from recently diverged taxa (Beardsley and Olmstead, 2002; Nie et al., 2006). However, our study also shows that even whole genome sequences might only imperfectly resolve phylogenetic relationships between closely related species. Therefore, genomic analysis of hybrid ancestry can be significantly strengthened by experimental crossing data such as the one conducted here. Exploration of reproductive isolation asymmetry across the plant kingdom through genome-skimming may allow for a better understanding of the origin of hybrid and allopolyploid species. Connecting the genomic and molecular underpinnings of reproductive asymmetry to the morphological and ecological consequences of asymmetric barriers will ultimately generate a better solution to the puzzle of allopolyploids’ evolutionary success.

## Acknowledgements

We thank H.E. O’Brien for comments and suggestions on haplotype analysis. We thank two anonymous reviewers for comments on a previous version of this manuscript. This work was supported in part by a UK Natural Environment Research Council grant (NERC, NE/J012645/1) and a Carnegie Trust Travel grant to MVM, and a Suzann Wilson Matthews Research Award to JRP.

